# Modulation of the E-cadherin in human cells infected *in vitro* with *Coxiella burnetii*

**DOI:** 10.1101/2022.12.08.519566

**Authors:** Ikram Omar Osman, Soraya Mezouar, Djamal Belhaouari-Brahim, Jean-Louis Mege, Christian Albert Devaux

**Affiliations:** Aix-Marseille Univ, IRD, APHM, MEPHI, Marseille, France; IHU-Méditerranée Infection, Marseille, France; CNRS, Marseille, France

**Author notes:** **Corresponding author:** Christian Devaux, PhD.

**Keywords:** *Coxiella burnetii*, E-cadherin, Soluble E-cadherin, β-catenin

## Abstract

High concentration of soluble E-cadherin (E-cad) was previously found in sera from Q fever patients. Here, BeWo cells which express a high concentration of E-cad were used as an *in vitro* model to investigate the expression and function of E-cad in response to infection by *Coxiella burnetii*, the etiological agent of Q fever. Infection of BeWo cells with *C. burnetii* leads to a decrease in the number of BeWo cells expressing E-cad at their membrane. A shedding of soluble E-cad was associated with the post-infection decrease of membrane-bound E-cad. The modulation of E-cad expression requires bacterial viability and was not found with heat-inactivated *C. burnetii*. Moreover, the intracytoplasmic cell concentration of β-catenin (β-cat), a ligand of E-cad, was reduced after bacterial infection, suggesting that the bacterium induces modulation of the E-cad/β-cat signaling pathway and *CDH1* and *CTNNB1* genes transcription. Finally, several genes operating the canonical Wnt-Frizzled/β-cat pathway were overexpressed in cells infected with *C. burnetii*. This was particularly evident with the highly virulent strain of *C. burnetii*, Guiana. Our data demonstrate that infection of BeWo cells by live *C. burnetii* modulates the E-cad/β-cat signaling pathway.

## Introduction

*Coxiella burnetii*, a known intracellular bacterium causing Query (Q) fever in humans, is most often transmitted to humans through products derived from infected animals[1], and primarily targets the lungs. The bacterium infects a large spectrum of susceptible cells, including myeloid cells[2,3], trophoblasts[4] and adipocytes[5]. Human primary infection remains asymptomatic in most people (around 60% of cases)[6]. In the other cases, symptoms develop within 2-6 weeks after *C. burnetii* exposure and mainly consist of hepatitis, endocarditis, pneumonia, vascular infection and lymphadenitis[7,8]. Although these symptoms usually resolve in a few weeks, in rare cases (less than 5%), the infection become persistent and progresses to chronic endocarditis[9]. Factors that may determine the severity of the disease remain mostly unknown[10–13], but considerable genomic heterogeneity was reported among *C. burnetii* strains[14–18], and gene deletion was associated with a higher strain virulence[19]. A role for the bacterial lipopolysaccharide (LPS) was also highlighted, since several *in vitro* passages of the virulent Nine Mile strain of *C. burnetii* generate an avirulent strain characterized by a truncated LPS with the absence of the terminal sugar-containing O-polysaccharide chain[19,20].

In the past decade, a diagnosis of Q fever with persistent focal infection was reported as an increased risk factor for B-cell lymphoma [21–23]. Overproduction of interleukin (IL)-10, a B-cell growth factor[24], is critical for sustaining *C. burnetii* replication[25], and might possibly favor lymphoma occurrence [22]. In peripheral blood mononuclear cells (PBMCs) of Q fever patients it was possible to characterize a transcriptional signature (overexpression of genes involved in the anti-apoptotic process or repression of pro-apoptotic pathways), that could be, in very rare cases, associated with the development of non-Hodgkin lymphoma (NHL)[26]. We also speculated that if *C. burnetii* infection could sometimes promote the development of NHL, the early steps towards NHL progression might be the decrease of E-cadherin (E-cad) surface expression on the CD20^+^ B-cells subpopulation of PBMCs (E-cad positive B-cells represents less than 1% of the circulating B lymphocytes) from *C. burnetii*-infected patients [27]. However, until now the molecular mechanisms that could account for the progression of the disease to lymphoma have remained elusive. Moreover, these results are controversial, and other studies of Q fever patients have not confirmed the link between Q fever and NHL[28]. A recent large analysis of the Dutch population incorporating the 2007-2010 Q fever outbreak (266 million people including 61424 diagnosed with NHL and 4310 persons diagnosed with acute Q fever), concluded to the lack of increased risk of NHL after Q fever[29].

We previously found that sE-cad is present in higher concentrations in sera of Q fever patients than healthy controls[27], and postulated that E-cad cleavage may be a step towards *C. burnetii-*invasion of the host during Q fever. Increasing evidence can be found in the literature indicating that cadherins may play a major role during the development of infectious diseases[30]. Among cadherins, the E-cadherin (E-cad), a 120 kDa adhesion cell-surface protein, is known for its role in cell-to-cell tight junctions ensuring integrity of epithelial barriers. E-cad also behaves as a signaling molecule through its intra-cytoplasmic tail that binds second messengers, thereby having roles in cell activation, cell division, and cell differentiation and/or invasion[31,32]. Besides its role in inter-cellular adhesion, E-cad behaves as a tumor suppressor[33–35] and plays a major role in cancer invasion/metastasis[36–38]. Cell reprogramming can be achieved by the release of sE-cad generated from the cleavage of the extracellular domain of E-cad[39]. E-cad is known to be a target for several bacterial sheddases, leading to the release of sE-cad[39,40].

Here we developed an *in vitro* cellular model with BeWo cells which express a high concentration of E-cad at their surface[41] and are susceptible to *C. burnetii* infection[4] to study the effect of *C. burnetii* infection on the expression of E-cad and evidenced that *C. burnetii* infection modulates expression of the E-cad/β-catenin signaling pathway by acting on *CDH1* and *CTNNB1* gene transcription, on cell-surface expression of E-cad, the release of sE-cad and the cytosolic pool of β-catenin.

## Materials and Methods

### Coxiella burnetii

*Coxiella burnetii* (Nine Mile strain RSA496, Guiana Strain Cb175) was grown as described[42]. L929 cells line were used for *in vitro* culture using a Minimum Eagle medium (MEM, Invitrogen, USA) supplemented with 4% fetal bovine serum (FBS, Invitrogen, USA) and 1% 2mM L-glutamine (Invitrogen, USA) and then incubated at 35°C in a 5% CO_2_ atmosphere. After three passages infected cells were sonicated, and the cell-free supernatants were centrifuged to harvest the bacteria, which were then washed and stored at −80°C until use. Gimenez staining and qPCR (using the *com-1* gene) were used to determine the concentration of bacteria in the sample.

### *In vitro* cell culture model

Blood samples (leucopacks) used in our study come from the French Blood Establishment (Etablissement français du sang, EFS), which carries out donor inclusions, informed consent and sample collection. Through a convention established between our laboratory and the EFS (No.7828), buffy coats were obtained from healthy blood donors. Peripheral blood mononuclear cells (PBMCs) were isolated after centrifugation on Ficoll cushions (MSL, Eurobio, France). Monocytes (5-15% of the total number of PBMCs) were obtained from PBMCs using magnetic beads coated with monoclonal Abs directed against CD14, according to the manufacturer’s instructions (Miltenyi Biotech, France). Monocytes-derived-macrophages (MDM) were obtained following the incubation of monocytes with 10% heat-inactivated human AB serum (MP Biomedicals, LLC, France) during a 3-day culture followed by 4 days of incubation with 10% FBS (Invitrogen, USA) as previously described[43]. Monocyte-derived-dendritic cells (moDC) were obtained by incubation of monocytes with 1 ng/mL of granulocytes-macrophages colony-stimulating factor (GM-CSF, R&D Systems, France) and Interleukin-4 (R&D Systems, USA) for 7 days as previously described[44]. Mast cells were obtained from PBMCs as previously described[45]. All cells were cultured in Roswell Park memorial institute (RPMI) 1640 medium (Gibco, Thermo Fisher, USA) supplemented with 10% FBS.

The human trophoblastic BeWo cell line was purchased from the American type culture collection (ATCC, CCL-98, Bethesda, USA) and cultured in a Dulbecco’s Modified Eagle Medium F-12 Nutrient Mixture (DMEM F-12, Invitrogen, USA) containing 10% FBS. The HeLa cell line (ATCC CCL-2), a human epithelial cell line derived from a cervical cancer, was cultured in DMEM supplemented with 10% FBS and 1% L-glutamine (Invitrogen, USA). MRC-5 (ATCC CCL-171), a human embryonic fibroblast cell line obtained from an aborted fetus, was cultured in MEM supplemented with 4% FBS and 1% L-glutamine. THP-1 (ATCC TIB-202), a human monocytic cell line derived from an acute monocytic leukemia patient, was maintained in RPMI 1640 medium supplemented with 10% FBS and 1% penicillin/streptomycin (Gibco) at 37°C and 5% CO_2_.

### Microarray and data analysis

The microarray was performed as previously described[26] with data submitted to NCBI’s Gene Expression Omnibus (GEO series accession number GSE112086). Briefly, RNAs were extracted using an RNeasy Mini Kit (QIAGEN SA, France) with a DNase I step, then labeled using Cyanin-3 CTP (Agilent Technologies) and hybridized on chips containing 45000 probes (4×44K Whole Human Genome). After hybridization, slides were washed and scanned with a pixel size of 5 μm using the DNA Microarray scanner G2505C. The raw data were extracted (Extraction Software 10.5.1) and processed using GeneSpring GX 14.9 software (Agilent Technologies). A selection filter of both statistical value p<0.01 (t-Student test) and an absolute value of fold change (FC) greater than 1.5 was used to identify differentially expressed genes. Modulated genes were analyzed using ClustVis software.

### Quantitative-Reverse Transcription Polymerase Chain Reaction (qRT-PCR)

BeWo cells (2×10^5^ cells/well) were cultured in flat-bottom 24-well plates for 12 h and were then infected with the live virulent *C. burnetii* Nine Mile (NMI) and Guiana (GuiI) strains or were exposed to heat-inactivated virulent *C. burnetii* NMI strains at a 100:1 bacterium-to-cell ratio, or stimulated with 100 ng/mL of lipopolysaccharide (LPS) from *E. Coli* (O55:B5; Sigma-Aldrich, USA) for 48 h at 37°C in a 5% CO_2_ atmosphere. RNAs were extracted as described above. The first-strand cDNA was obtained using oligo(dT) primers and Moloney murine leukemia virus-reverse transcriptase (MMLV-RT kit; Life Technologies, USA), using 100 ng of purified RNA. The qPCR experiments were performed using specific oligonucleotide primers and hot-start polymerase (SYBR Green Fast Master Mix; Roche Diagnostics, Germany). The amplification cycles were performed using a C1000 Touch Thermal cycler (Biorad, USA). Specific primers used in this study are listed in **Table 1** and the results were normalized using the housekeeping gene β-actin (*ACTB*) (Fwd: 5’ CAT GCC ATC CTG CGT CTG GA 3’; Rev: 5’ CCG TGG CCA TCT CTT GCT CG 3’) and expressed as the relative expression (2^-ΔCT^) where ΔCt = Ct_Target_ − Ct_Actin_, and as the fold change (2^-ΔΔCT^), where ΔΔCt = ΔCt_stimulated_ - ΔCt_unstimulated_ as previously described[46]. Ct values were defined as the number of cycles for which fluorescence signals were detected.

**Table 1.**
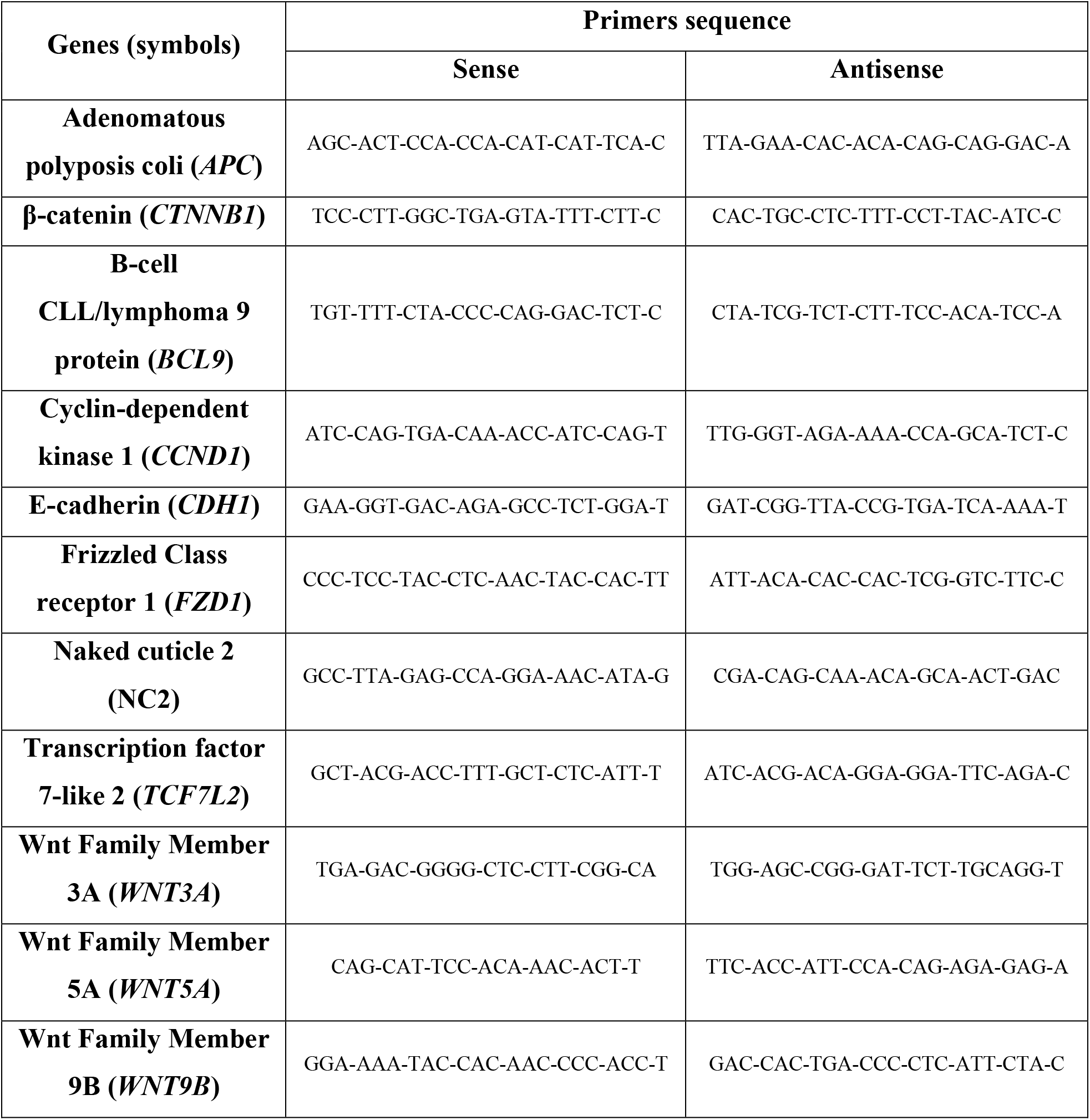
Specific primers used for q-RTPCR assays.

### E-cadherin protein quantification

BeWo cells (5×10^5^ cells/well) were cultured in flat-bottom 12-well plates for 12 h and were then infected with live *C. burnetii* NMI strains or exposed to heat-inactivated *C. burnetii* strains at a 50:1 bacterium-to-cell ratio or stimulated with 100 ng/mL of LPS for 4, 24 and 48 h at 37°C in a 5% CO_2_ atmosphere.

For each kinetics, the culture supernatants were collected, centrifuged at 1000 g for 10 min and stored at −20°C until use. The quantity of sE-cad in the supernatants was determined using a specific immunoassay (DCADEO, R&D Systems, USA) according to the manufacturer’s instructions. The minimal detectable concentration of human sE-cad was 0.313 ng/mL.

Protein quantification was evaluated using a western blot assay. Cells were washed with ice cold phosphate buffered saline (PBS 1X) and lysed using 1X RIPA buffer (100 mM Tris-HCl pH 7.5, 750 mM NaCl, 5 mM EDTA, 5% IGEPAL, 0.5% sodium dodecyl sulfate (SDS), and 2.5% Na deoxycholate) supplemented with both a protease and phosphatase cocktail inhibitor (Roche, Germany). Ten μg of protein were loaded onto 10% SDS polyacrylamide gels. After transfer into a nitrocellulose membrane, the blots were incubated overnight at 4°C with a saturation solution (5% fat free milk (FFM)-1X PBS - 0.3% Tween 20). Blots were then incubated with mouse anti-human E-cad ectodomain mAb (1:5000) (HECD-131700, Invitrogen, USA) for 2 h, followed by an incubation with mouse anti-human E-cad cytoplasmic domain mAb (4A2C7, Life Technologies). In some experiments, β-catenin was detected using an anti-human β-catenin antibody (1:1000) (Agilent Dako, USA). After three washes in 1X PBS – 0.3% Tween 20, the blots were incubated with horseradish peroxidase-conjugated sheep anti-mouse IgG (1:10000) for 2 h at room temperature. β-actin expression was measured using an anti-human β-actin horseradish peroxidase-conjugated mAb (1:25000, Life Technologies, USA) as the loading control. The proteins were revealed using an ECL western blotting substrate (Promega, USA) and images were digitized using a Fusion FX (Vilber Lourmat, France). The density of the bands was measured using Image J v 1.8.0.

### Confocal immunofluorescence assay

BeWo cells were cultured on sterile coverslips in 24-well plates at an initial concentration of 2×10^5^ cells/well and were then infected with live virulent *C. burnetii* NMI strains or exposed to heat-inactivated virulent *C. burnetii* NMI strain at a 50:1 bacterium-to-cell ratio or stimulated with 100 ng/mL LPS for 24 h at 37°C in a 5% CO_2_ atmosphere. After fixation with 4% paraformaldehyde, cells were permeabilized with 0.1% Triton X-100 for 3 min and saturated with 3% BSA – 0.1% Tween 20 - PBS for 30 min at room temperature. Fixed cells were first incubated with a mouse anti-E-cad cytoplasmic domain mAb (1:1000) (4A2C7, Life Technologies, USA) for 1 h at room temperature and then with 1:500 rabbit polyclonal anti-*C. burnetii* antiserum. After washing, cells were incubated for 30 min at room temperature with a mix (1:1000) of goat anti-rabbit IgG secondary antibody (Alexa Fluor 647, Life Technologies, USA) and goat anti-mouse IgG secondary antibody (Alexa Fluor-555, Life Technologies, USA). 4’,6’-diamino-2-fenil-indol (1:25000) (DAPI, Life Technologies, USA) and Phalloidin (1:500) (Alexa-488, OZYME, France) were used for staining the nucleus and the filamentous actin, respectively. Labelled cells were analyzed using laser scanning confocal microscopy and pictures were acquired using a confocal microscope (Zeiss LSM 800) with a 63X/1.4 oil objective, an electronic magnification of 0.7 and a resolution of 1014_1014 pixels.

### Flow cytometry assay

BeWo cells (1×10^6^ cells/well) were cultured in flat-bottom 6-well plates and were then infected with a live virulent *C. burnetii* NMI strain at a 50:1 bacterium-to-cell ratio or stimulated with 100 ng/mL LPS for 24, 48 and 72 h at 37°C in a 5% CO_2_ atmosphere. Cells were incubated with 2 mM EDTA in PBS for 15 min at 4°C to release the adherent cells from the culture plate. After centrifugation at 500 g for 5 min, the pellet was suspended in a FACS buffer (2 mM EDTA, 10% FBS in PBS) and then incubated for 30 min with a mouse anti-human E-cad ectodomain antibody (HECD-131700, 1:1000). After two washes, cells were incubated with a goat anti-mouse IgG Alexa Fluor-555 secondary antibody. Fluorescence intensity was measured using a Canto II cytofluorometer (Becton Dickinson/Biosciences, France) and the results were analyzed using FlowJo V10.7.2 software (Becton Dickinson, USA).

### Electron microscopy analysis

BeWo cells (1×10^6^ cells/well) were cultured in flat-bottom 6-well plates and were then infected with live virulent *C. burnetii* NMI and GuiI strains at a 100:1 bacterium-to-cell ratio at 37°C in a 5% CO_2_ atmosphere. After 24 h of infection, the cells were fixed with 2.5% glutaraldehyde in 0.1 M cacodylate buffer for five hours at 4°C. Resin embedding was microwave-assisted with a Biowave Pro+ (Pelco, Fresno, CA, USA). After washing twice with a mixture of 0.2-M saccharose/0.1-M sodium cacodylate and once with distilled water, samples were progressively dehydrated by successive baths in 50%, 70% and 96% ethanol. Substitution with LR–White resin (medium grade; Polysciences, Warrington, PA, USA) was achieved by incubations with 25% to 100% LR–White resin and samples were placed in a polymerization chamber for 72 h at 60°C. All solutions used above were 0.2-μm filtered. After curing, the resin blocks were manually trimmed with a razor blade and dish bottoms were detached from cell monolayers by cold shock via immersion in liquid nitrogen for 20 seconds. Resin blocks were placed in a UC7 ultramicrotome (Leica), trimmed to pyramids, and ultrathin 100 nm sections were cut and placed on HR25 300 Mesh Copper/Rhodium grids (TAAB, Aldermaston, England). Sections were contrasted with uranyl acetate and lead citrate. Grids were attached with double sided tape to a glass slide and platinum-coated at 10 mA for 20 seconds with a MC1000 sputter coater (Hitachi High-Technologies, Japan). Electron micrographs were obtained on a SU5000 SEM (Hitachi High-Technologies, Japan) operated in high-vacuum at 10 kV accelerating voltage and observation mode (spot size 30) with BSE detector.

### Statistical analysis

The statistical analyses of the data were performed using GraphPad-Prism software (version 6.0). The results are presented as the ± standard error of the mean (SEM). The Mann-Whitney U test was used for flow cytometry, the two-way ANOVA test for E-cadherin quantification and transcriptional analysis, and a nonparametric Kruskal-Wallis with Dunn’s multiple comparison test for group comparison. A p value <0.05 was considered statistically significant.

## Results

### Selection of the BeWo cell line as an appropriate cellular model to study E-cad expression in cells infected with *C. burnetii*

It was previously reported that many cell types, including myeloid cells[47,48], trophoblasts[4,49], lung cells[50,51], and adipocytes[5] are susceptible to *C. burnetii* infection. To conduct our study, it was first necessary to select a cellular model with high expression of E-cad. To this end, we evaluated E-cad expression in different cell lines, including HeLa (human cervical cancer cell line), THP-1 (human monocytic cell line), MRC-5 (human fetal lung fibroblasts), BeWo (human placenta choriocarcinoma cell line), and primary monocytes from healthy donors. As shown in **Figure 1A**, using a western blot assay, BeWo cells were found to express the highest concentration of E-cad and HeLa cells ranked second. Additionally, we also confirmed the highest membrane expression of E-cad on BeWo cells by confocal microscopy (**Figure 1B**). This finding was consistent with the reports from the literature on the common use of BeWo cells as a model to study E-cad expression[52,53]. Moreover, scanning electron microscopy of *C. burnetii*-infected BeWo cells (**Figure 1C** and **Figure S1**) showed the presence of several vacuoles, either empty or containing electron-dense circular structures, with diameters ranging from 250 nm to 700 nm dispersed in the cell cytoplasm, corresponding to the morphology and size of *C. burnetii* as previously described[54]. Therefore, the BeWo cell line was chosen for further investigation because it has both sensitivity to *C. burnetii* infection and high expression of E-cad.

**Figure 1.**
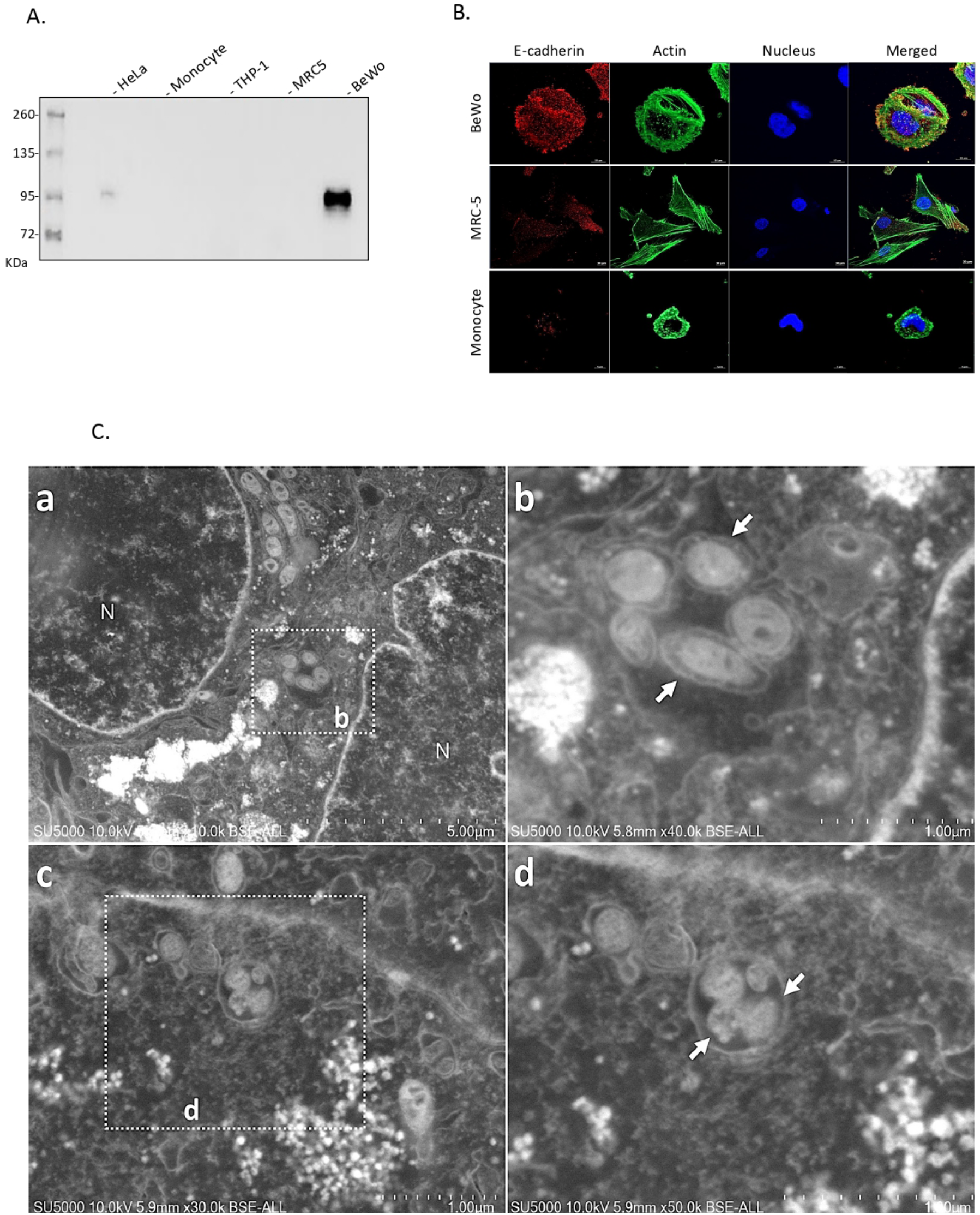
Selection of a cellular *in vitro* model to study E-cad modulation. **(A)** Immunoblot detection of E-cad expression in protein lysates of various cell types including HeLa, THP-1, MRC-5, and the BeWo cell line and primary human monocytes **(B)** Cellular localization/distribution of E-cad protein (red), actin (green) and nucleus (blue) in three different cell types (BeWo cells, MRC-5 cells, monocytes). Images were acquired using a confocal microscope (Zeiss LSM 800) with a 63X/1.4 oil objective (Scale bar: 10 μm). (**C**) Scanning electron microscopy images of the ultra-thin section of BeWo cells infected with virulent *Coxiella burnetii* Nine Mile (**a**) or *Coxiella burnetii* Guiana strain (**c**). **(b)** and (**d**) respectively high magnification of the boxed region in **(a)** and **(c)** show a vacuole-contained *Coxiella burnetii*-like bacteria (white arrows).

### *C. burnetii* infection modulates cell surface expression of E-cad

The level of E-cad protein expression at the cell surface of BeWo cells infected or not by *C. burnetii* NMI was evaluated by flow cytometry, targeting the ectodomain portion of E-cad. Twenty-four hours post-*C. burnetii* infection, the percentage of BeWo cells expressing membrane-bound E-cad decreased significantly compared to LPS stimulated- and unstimulated-cells (p<0.05 and p<0.01, respectively) used as controls (**Figure 2A and 2B**). Also, we found a significant decrease in the mean intensity fluorescence signal of the E-cad in BeWo cells infected with *C. burnetii* compared to LPS-stimulated and unstimulated cells (p<0.05 and p<0.01, respectively) (**Figure 2C**). Then, we studied whether bacterial antigen/cell contact was sufficient to trigger E-cad modulation, or whether infection was absolutely required to observe a change in E-cad expression. Thus, we investigated by confocal microscopy E-cad expression on BeWo cells after 4 hours and 24 hours of infection with live *C. burnetii*, while controls consisted of exposure to heat-inactivated bacteria or LPS. Although for unstimulated BeWo cells a high level of fluorescence and homogeneity of E-cad distribution was observed, this result was not limited to this experimental condition. The same expression profiles were observed for BeWo cells after *E. coli* LPS stimulation or heat-inactivated *C. burnetii* bacterium stimulation (**Figure 2D**). Interestingly, live *C. burnetii*-infected BeWo cells presented a significant decrease of E-cad expression and also a short intracellular colocalization of the E-cad fluorescence signal with the fluorescence signal associated with *C. burnetii* at early stage of infection (**Figure S2**). In contrast, a low E-cad fluorescence signal was observed in infection (live *C. burnetii*-infected BeWo cells). This decrease of E-cad cell-surface expression was accompanied by an actin rearrangement over time (as shown at 24 hours) which was faster when the cells where infected with the virulent Guiana strain (**Figure S3**). These results indicate that membrane-bound E-cad expression is highly modulated during active *C. burnetii* infection of BeWo cells, while *C. burnetii* bacterial antigens are not sufficient to trigger a massive E-cad modulation.

**Figure 2.**
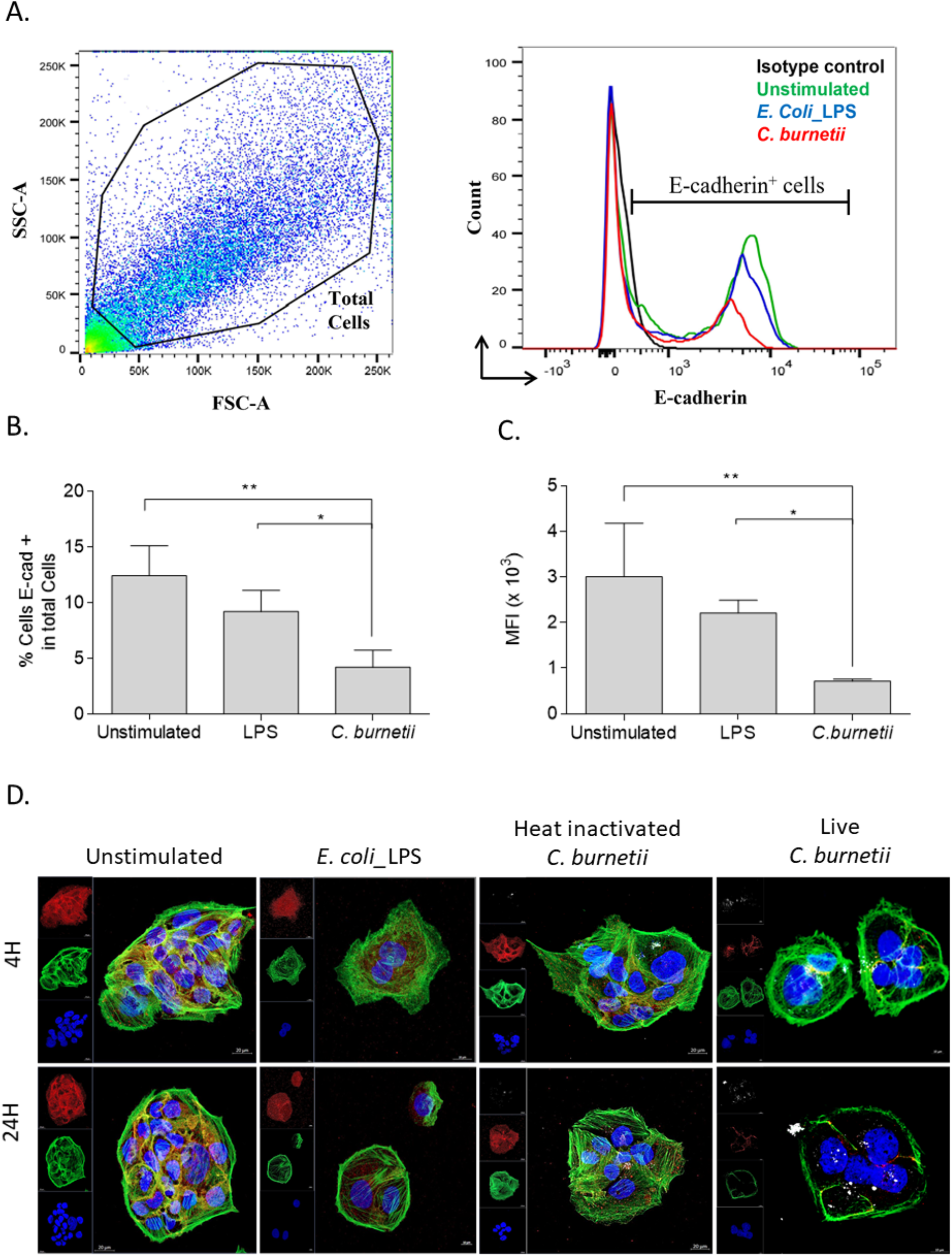
Modulation of E-cad expression in BeWo cells either exposed to live *C. burnetii* or *E. coli* LPS. **(A)** Flow cytometry analysis of E-cad expression at the surface of BeWo cells. Cells were incubated with live *C. burnetii* Nine Mile strain or were stimulated with *E. coli* LPS for 24 hours (n = 3). BeWo cells maintained in cultured medium without additive (unstimulated) were used as control. The two panels show the gating parameter chosen and the quantification of E-cad, respectively. **(B)** Histogram indicating the percent of cells expressing E-cad. **(C)** Histogram representing the mean fluorescence intensity (MFI) under the different experimental conditions. The non-parametric Mann-Whitney test was used for statistical analysis of all data. The p value < 0.05: symbol *; p value <0.01: symbol **. **(D)** The panel presents *C. burnetii* (live or heat-inactivated bacterium) exposed cells versus *E. coli* LPS stimulated cells and evaluation of fluorescence corresponding to the E-cad (red), *C. burnetii* (white), actin (green), and the nucleus of the cells (blue). Images were acquired using a confocal microscope (Zeiss LSM 800) with a 63X/1.4 oil objective (scale bar: 20 μm).

### Release of sE-cad by *C. burnetii*-infected BeWo cells and modulation of *CDH1/*E-cad and *CTNNB1*/β-catenin gene transcription

Since we previously reported that Q fever patients have high expression of serum sE-cad[27], we aimed at investigating whether the decreased expression of membrane-bound E-cad in *C. burnetii*-infected BeWo cells could be associated with a release of sE-cad after E-cad proteolysis. As shown in **Figure 3A**, we measured progressive basal accumulation of sE-cad in the cell culture medium over time (at 24 hours and 48 hours). Interestingly, sE-cad release was significantly higher (p <0.001) over time in infection conditions compared to control conditions (unstimulated and LPS-stimulated BeWo cells). A further immunoblot detection of the products synthesized in *C. burnetii-* infected cells evidenced two components: the integral 120 kDa E-cad, and a protein of about 60 kDa, a proteolytic product of E-cad (**Figure 3C**). In contrast, only the 120 kDa E-cad was found in the control experiments. It is worth noting that the expression of β-catenin, an intracellular ligand of E-cad, was strongly decreased in BeWo cells under live *C. burnetii*-infection conditions. This was further confirmed using a densitometry scanning of the band that showed a significant (p<0.05) decrease of β-catenin in *C. burnetii*-infected BeWo cells.

**Figure 3.**
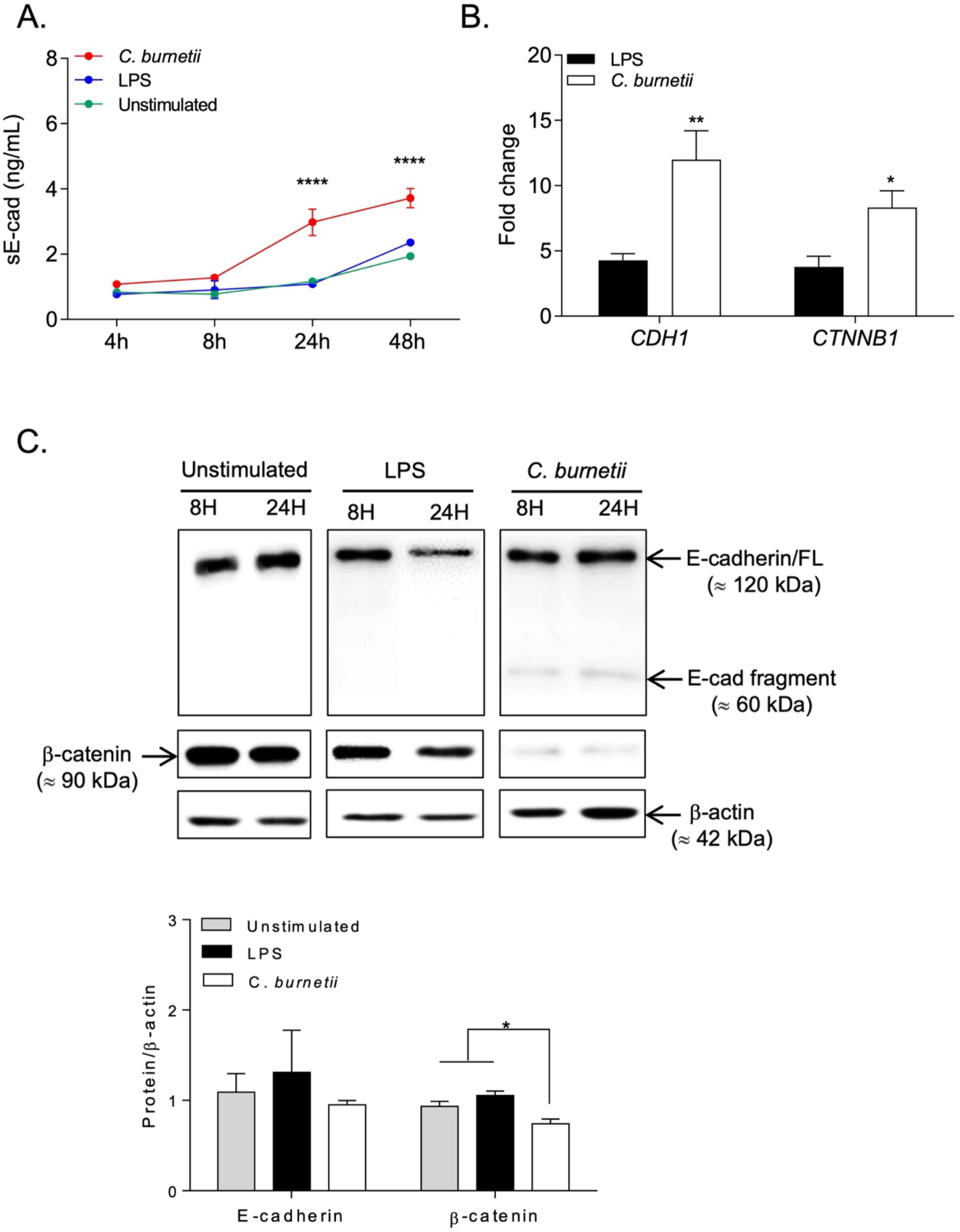
Impact of *C. burnetii* infection of BeWo cells on their E-cad expression. BeWo cells were incubated with live *Coxiella burnetii* Nine Mile strain for 4, 8, 24, and 48 hours (n=3). **(A)** ELISA quantification of sE-cad released in the supernatant of BeWo cells under different experimental conditions (unstimulated, *E. coli* LPS-stimulated, *C. burnetii* Nine Mile strain-infected), at different times of experiment. **(B)** qRT-PCR analysis of *CDH1*/E-cadherin and *CTNNB1*/β-catenin mRNAs expression in BeWo cells exposed *in vitro* during 48 hours to *C. burnetii* Nine Mile strain or the *E. coli* LPS. Expression level of investigated genes was illustrated as ± SEM of the fold change (FC = 2^-ΔΔCt^, where ΔΔCt = [(Ct_target_ – Ct_actin_) stimulated] – [(Ct_target_ – Ct_actin_) unstimulated] and p values was evaluated according to an ordinary Two-way Anova test). **(C)** Blots were spliced after protein transfer and then were probed with antibodies. Upper panel: Immunoblot detection of E-cad (upper blot), β-catenin (middle blot) and β-actin (lower blot) expression in total protein lysates of BeWo cells cultured under the three different experimental conditions described above (unstimulated, LPS-stimulated, *C. burnetii*-infected), at 8 and 24 hours post treatment. The histogram shown in the lower panel is a quantification of E-cad and β-catenin expression (n=3) with respect to β-actin. The p value < 0.05: symbol *; p value <0.01: symbol **

In accord with the literature regarding the E-cad/β-catenin signaling pathway, our results suggest a translocation of β-catenin into the cell nucleus, associated with a modulation of *CDH1*/E-cad gene expression. To confirm this hypothesis, the expression of *CDH1*/E-cad and *CTNNB1*/β-catenin gene transcription was analyzed by qRT-PCR. Both *CDH1* and *CTNNB1* genes were found significantly overexpressed (p<0.01 and p<0.05, respectively) in BeWo cells with *C. burnetii* infection compared to cells exposed to *E. coli*-LPS (**Figure 3B**). These results indicate that *C. burnetii* infection of BeWo cells induces a shedding of the E-cad protein ectodomain at an early stage of the infection (detectable at 24 hours post-infection) and suggest that sE-cad shedding is associated with the nuclear translocation of β-catenin and subsequent up-regulation of *CDH1* and *CTNNB1* gene transcription through a feedback control loop.

### *C. burnetii* modulates expression of genes from both the E-cad/β-catenin pathway and the Wnt/frizzled/β-catenin pathway in BeWo cells

To further demonstrate that the overexpression of *CDH1* and *CTNNB1* genes found in *C. burnetii* infection of BeWo cells is not strain-dependent, BeWo cells were exposed for 48 hours to either the live Nine Mile strain of *C. burnetii* or the more virulent Guiana (Cb175) laboratory strains of *C. burnetii*. Heat-inactivated *C. burnetii* (Nine Mile strain) and *E. coli* LPS were used as controls.

As shown in **Figure 4A and 4B**, a significant overexpression of *CDH1* and *CTNNB1* genes was found in BeWo cells infected with each of the two live strains of *C. burnetii*, extending our earlier observations, illustrated in **Figure 3C**. In contrast, when BeWo cells were exposed to heat-inactivated bacteria, *CTNNB1* gene expression was further increased and *CDH1* was significantly decreased. These data suggest that the interaction of bacterial antigens with BeWo cells is sufficient to trigger signals leading to *CTNNB1* gene transcription, while induction of *CDH1* requires infection.

**Figure 4.**
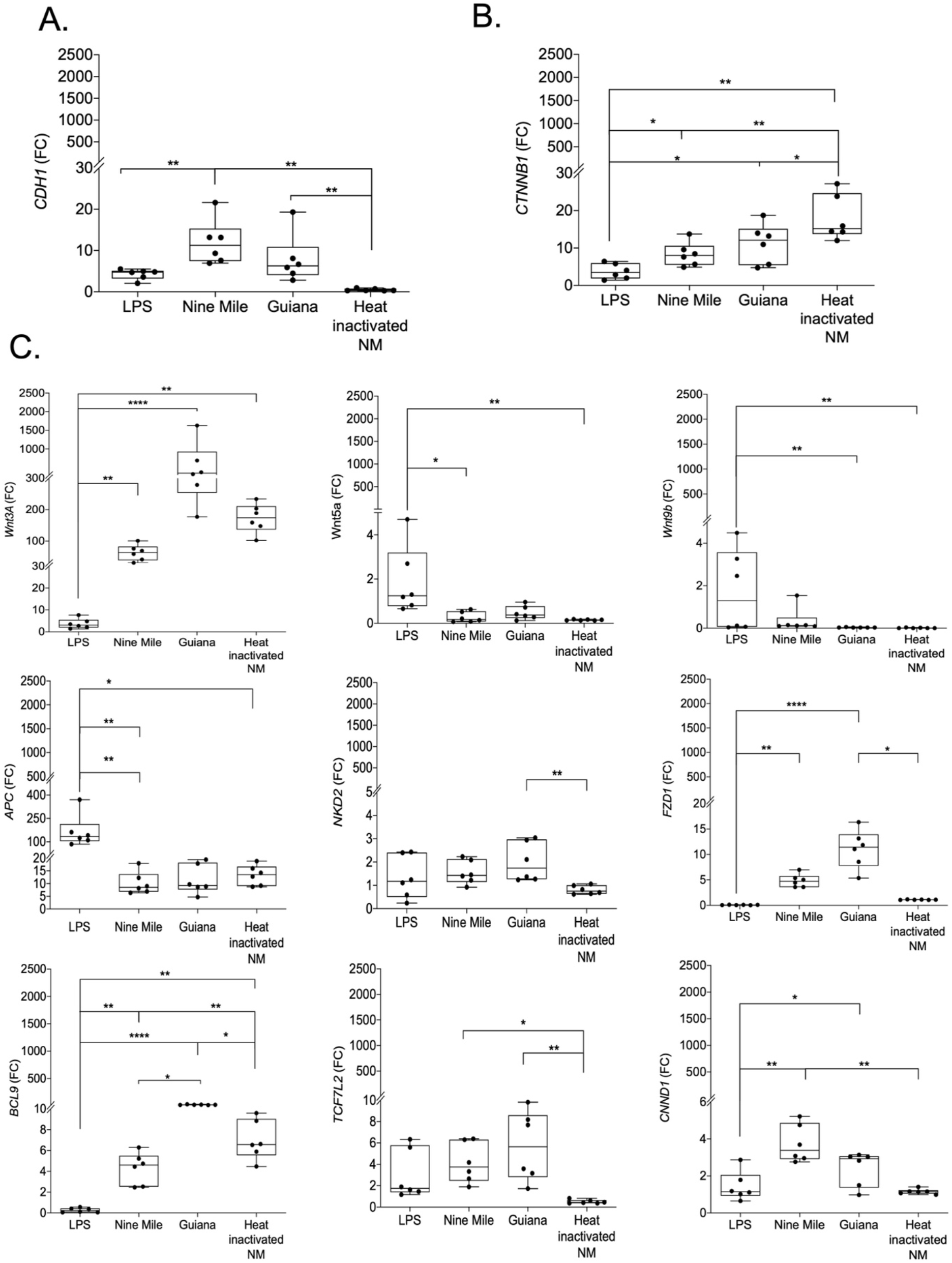
Impact of *C. burnetii* (Nine Mile strain or Guiana strain) infection of BeWo cells on the expression of *CDH1*/E-cad, *CTNNB1*/β-catenin and other genes involved in the Wnt/frizzled/β-catenin pathways. BeWo cells were exposed in *vitro* for 48 h to *C. burnetii* Nine Mile strain or Guiana (Cb 175) strain (live and heat-inactivated strain) or were stimulated with *E. coli* LPS. The expression (n=6) of *CDH1*/E-cad **(A)**, *CTNNB1*/β-catenin **(B)** and other genes involved in the Wnt/frizzled/β-catenin pathways **(C to K)**, was investigated by qRT-PCR after normalization with the housekeeping actin gene as endogenous control. Expression level of investigated genes was illustrated as ± SEM of the fold change (FC = 2^-ΔΔCt^, where ΔΔCt = [(Ct_target_ – Ct_actin_) stimulated] – [(Ct_target_ – Ct_actin_) unstimulated] and compared using the Kruskal-Wallis test. The p value < 0.05: symbol *; p value <0.01: symbol **; p value < 0.001: symbol ***; p value <0.0001: symbol ****

We then explored the expression of several other genes known to encode proteins linked to the E-cad/β-catenin and Wnt/frizzled/β-catenin axis, including *WNT3A, WNT5A, WNT9b, FZD1, APC, NKD2, BCL9, TCF7L2* and *CNND1*. As shown in **Figure 4C**, most of these genes were overexpressed in *C. burnetii-*infected BeWo cells, some of which (e.g., *BCL9, Wnt3A*) being also overexpressed when BeWo cells were exposed to heat-inactivated-bacteria infected cells. Notably, *BCL9, WntA3*, and *FDZ1* genes were significantly overexpressed where cells were infected with the live and more virulent Guiana strain of *C. burnetii* compared to infection with the live Nine Mile strain. We also observed that the gene coding for the tumor suppressor *APC* was significantly overexpressed after BeWo cell exposure to *E. coli* LPS. It is worth noting that the *CCND1* gene coding for cyclin D1 kinase, a cell cycle activator and oncogenic protein, was significantly over-transcribed in cells infected with live *C. burnetii*, notably in cells infected with the Nine Mile strain. The *CCND1* gene is much less expressed with heat-inactivated bacteria, suggesting that activation of this gene requires infection of cells by live bacteria. In addition, we also used a micro-array technology to compare the expression of a set of 45000 genes in BeWo cells, infected or not by *C. burnetii*. This approach indicated that 12189 genes were down-modulated after live *C. burnetii* infection of BeWo cells, while 6492 genes were found up-regulated, among which genes such as *CDH1, CCND1, and FZD1* were found highly expressed compared to their transcription levels in uninfected BeWo cells (**Figure S4**).

The transcriptomic results illustrated in Figure 4 were subjected to a principal component analysis (PCA) and a hierarchical clustering heatmap was regenerated. This analysis highlighted distinct gene expression profiles (**Figure 5**), the first one corresponding to live *C. burnetii* strain infectious conditions, with slight differences between live *C. burnetii* strains, while other profiles corresponded to unstimulated cells, LPS stimulated cells and cells exposed to heat-inactivated *C. burnetii*. Genes activated by the live *C. burnetii* strain infection cluster together and show a mirrored gene expression compared to other conditions. These results indicate that live *C. burnetii* specifically induce *CDH1, CCND1, CTNNB1* and *BCL9* gene overexpression in BeWo cells. However, for induction of the BCL9 gene, infection is not required, and transcription can be stimulated by simply exposing the cells to *C. burnetii* bacterial antigens.

**Figure 5.**
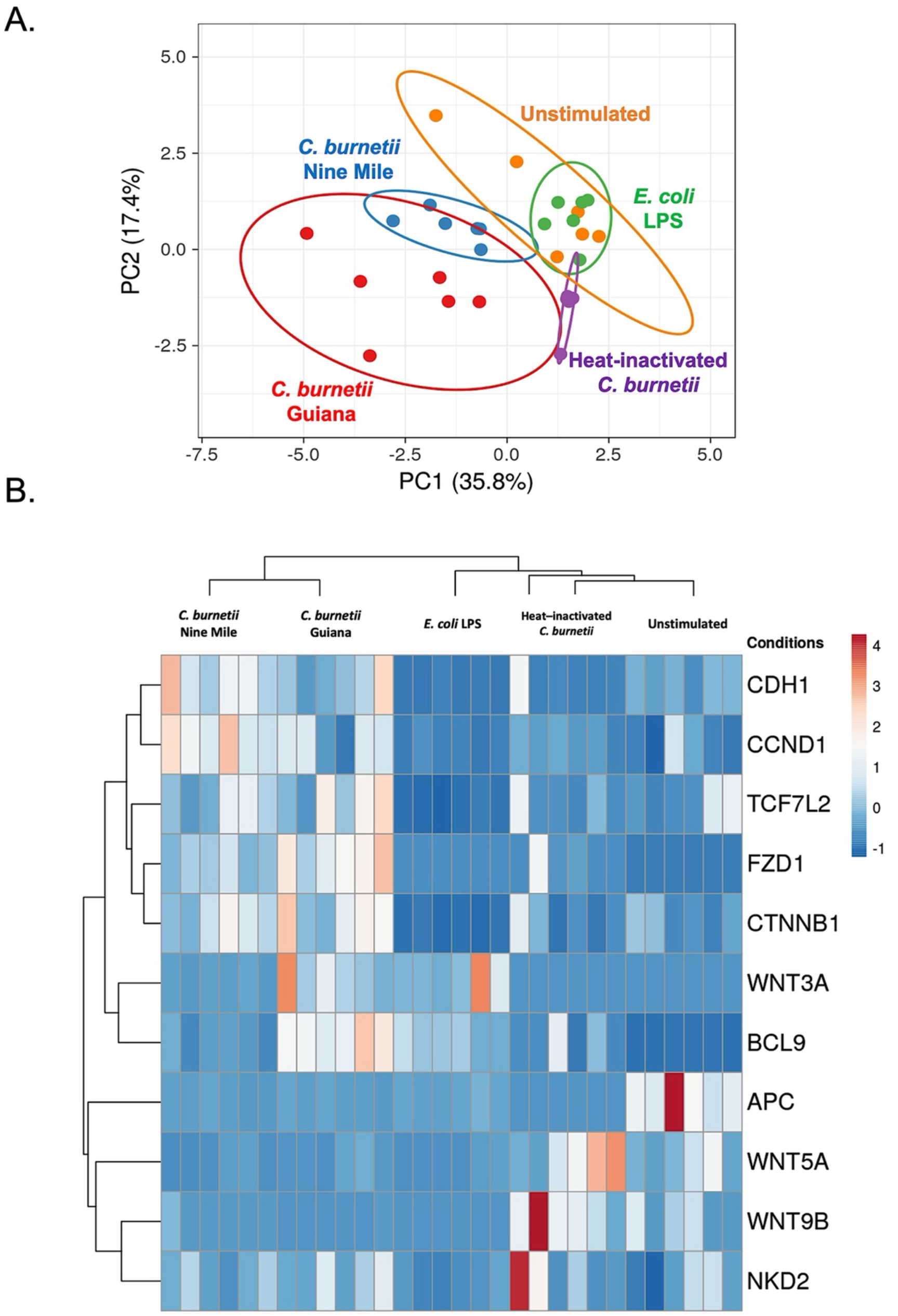
Principal component analysis and hierarchical clustering. BeWo cells were infected or not with different strains of *C. burnetii* (live or heat inactivated) for 48 h. The transcripts of several genes were quantified by qRT-PCR. Results (n=6) were expressed as RE = 2^-ΔCT^, where Δct = (Ct_target_ – Ct_actin_). Principal component analysis **(A)** and hierarchical clustering **(B)** were performed to show the distribution of the different groups of infected/stimulated BeWo cells and the modulation of expression of different genes, including *CDH1*/E-cad, *CTNNB1*/β-catenin and other genes involved in the Wnt/frizzled/β-catenin pathways, such as WNT3A, WNT5A, WNT9B.

## Discussion

We report here the first experimental evidence that differential gene expression analysis shows modulation of the E-cadherin/β-catenin pathway signature during *C. burnetii* infection *in vitro* and that this modulation results in disruption of E-cad expression on the cell surface. As a strictly intracellular bacterium, and in order to establish a replication niche, *C. burnetii* forms intimate interactions with its hosts, which aid in avoidance of the host immune response. In addition to the immune response, *C. burnetii* must overcome another central host defense mechanism, apoptosis. The molecular crosstalk between the bacterium and the host affects the induction of apoptosis, thereby prolonging its survival.

We previously reported that activation of protein tyrosine kinases (PTK) by *C. burnetii* in TPH-1 monocytes reflects *C. burnetii* virulence, since avirulent variants were unable to stimulate PTK[55]. Additionally, modulation of cell signaling following *in vitro C. burnetii* infection of the TPH-1 human macrophage-like cell line was also reported by Voth and Heinzen[56], who found that infection directs the sustained activation of host pro-survival kinases Akt and Erk1/2, necessary for anti-apoptotic activity that can conduct the development of certain lymphomas. Menelotte et al.[22] were able to link the development of B-cell non-Hodgkin lymphoma to *C. burnetii* infection.

Despite intensive work to identify the bacterial compounds supporting *C. burnetii* replication and virulence[11–13], the molecular mechanisms involved in *C. burnetii*-induced Q fever and possibly associated NHL remain largely unknown. In more recent investigations[26] and working on Q fever patient samples, we found that specific genes involved in anti-apoptotic processes (e.g., *BCL-2*) were highly expressed, whereas pro-apoptotic genes were repressed in PBMCs from patients with *C. burnetii-*associated NHL, supporting possible involvement of the corresponding proteins in lymphomagenesis. However, these results remain controversial since other studies have not confirmed the link between Q fever and NHL[28],[29].

The aim of the present study was to investigate *in vitro* the ability of *C. burnetii* to induce a modulation of the E-cad/β-cat axis during infection. For this purpose, BeWo cells, which are susceptible to *C. burnetii* infection, were used for their high expression of E-cad at the cell membrane. As a result, we were able to demonstrate that the presence of *C. burnetii* in the cells triggers overexpression of most of the genes of the signaling pathway, in particular *CDH1* genes coding for the E-cad protein. Indeed, E-cad is known as a tumor suppressor trans-membrane protein that inhibits β-cat activity by sequestrating it into the cell cytoplasm through interaction *via* the E-cad cytoplasmic domain. In addition, cleavage of the E-cad protein by proteases (known as sheddases) and excretion of a soluble fraction of E-cadherin (sE-cad) can contribute to cellular transformation through the loss of cell-to-cell adhesion[57]. This is considered to be a triggering event, promoting translocation of β-cat to the nucleus, where it forms complexes with BCL9 and T-cell factor (TCF), leading to transactivation of cell cycle genes such as cyclin kinase *D1/CCND1* and increased cell proliferation[58,59]. Moreover, the generation of sE-cad through sheddases and the subsequent nuclear re-localization of β-cat may be a critical indicator for cancer development. The induction of carcinogenic cascades via the E-cad/β-cat axis was reported in gastric adenocarcinoma associated with *Helicobacter pylori* infection, in which the bacterium activates calpain sheddase[60], or in myeloid-cell dependent distal colon tumorigenesis associated with *Bacteroides fragilis* toxin, a known bacterial sheddase[61,62]. We recently reported[27] that E-cad is cleaved in cells infected by *C. burnetii*. We demonstrated that, in addition to modulating *CDH1* expression at the transcriptomic level, infection with *C. burnetii* leads to disruption of E-cad at the cellular surface of BeWo cells. Under infection conditions, we observed a reduction of E-cad expression at the cell surface that is likely due to cleavage of the extracellular fraction of the protein, as its concentration in the supernatant is continuously increasing over time and the *CDH1* gene is overexpressed. In BeWo cells infected with live *C. burnetii*, we observed a remodeling of the actin filament, which triggers a morphological change. This particular effect was reported for the first time in the study of Meconi et al.[63,55], in which they showed that virulent *C. burnetii* stimulated morphological changes in human monocytes via the reorganization of the actin cytoskeleton. As E-cad is linked to the actin cytoskeleton through β-cat and α-cat, a rearrangement of the actin bundle can modify the adhesive function of E-cad and play an active role in the migratory activity of carcinoma cells[64]. In the case of *C. burnetii* infection, the bacterium, through the disruption of E-cad, may exploit the actin cytoskeleton to modulate its internalization by host cells. An interesting observation that emerged from this study concerns the decreased amount of β-cat protein in the cells, even though it is overexpressed at the transcriptomic level in cells infected with the live bacterium. As β-cat is part of the canonical Wnt-Frizzled/ β-cat pathway, we investigated most of the gene operating in the axis and as a result the expression of the genes *WNT3A* (encoding the extracellular Wnt molecules that bind Frizzled/low-density lipoprotein-receptor related proteins (LPR) complex and act on the canonical Wnt signaling pathway), *FZD1*/Frizzled (Wnt receptor), *NKD2* (Wnt pathway regulator protein) were significantly overexpressed in cells infected with live bacterium, mainly those infected with the more virulent Guiana strain (Cb175). Interestingly, it has been shown that the activation of the Wnt/ Frizzled//β-catenin pathway led to altered expression of genes involved in cell cycle regulation and apoptosis in normal and leukemic B-cell progenitors[65]. In a previous publication, an overexpression of transcription factor TCF7L2 and BCL9 was found, which is known to promote tumor progression by conferring enhanced proliferation, metastatic, and angiogenic properties to cancer cells[66]. This may indicate that in the extremely rare cases of patients with *C. burnetii*-associated NHL, a possible activation signal leads to lymphoma initiation. Furthermore, we also note that the expression of the tumor suppressor adenomatous polyposis coli (APC) is repressed in infected cells compared to the other conditions, suggesting that *C. burnetii* promotes the canonical Wnt signaling pathway. Similar results were obtained from the microarray analysis of the Wnt/Frizzled/β-catenin pathways in BeWo cells gene expression. Obviously, we are well aware of having used in this study a particular cell model, a human trophoblastic cell line, and that under these conditions there is a risk to overstate the biological significance of our data with the aim of explaining certain aspects of the pathophysiology of Q fever. In fact, B cell analysis, which would have been more physiologically relevant, was not possible because the CD20+E-Cad+ cells represent less than 1% of circulating B cells, as we published in a previous article[27]. However, this BeWo model is very useful for studying the link between infection by *C. burnetii* and modulation of the expression of E-cad and provided a possible molecular basis for explaining the increase in sE-cad concentrations in sera of Q fever patients [27].

In conclusion, this work provides for the first time evidence of the disruption of E-cad both at the transcriptomic and cell surface levels by *C. burnetii*, opening a new avenue of research in understanding the pathophysiology of Q fever. We hypothesize that the cleavage of E-cad by *C. burnetii* could be a means to ensure its transmigration in order to modulate the reactivity of immune cells and thus facilitate *C. burnetii* progression towards the target organs. Upon further investigation, we found that the cleavage of E-cad was mediated by a functional HtrA sheddase encoded in the *C. burnetii* genome[67]. Hence, the release of sE-cadherin together with the overexpression of Wnt/Frizzled/β-catenin may participate in the molecular crosstalk that occurs in the lymph node microenvironment during persistent Q fever and, in very rare cases, triggering a pro-carcinogenic program. Finally, it can be hypothesized that the decrease in the surface expression of E-cad induced by *C. burnetii* influences the replication of the bacterium or its virulence.

## Acknowledgments

We thank Prof. Jean Christophe LAGIER and Didier RAOULT for stimulating discussions and unwavering support.

## Funding

This research received no external funding.

## Supplementary data

**Figure S1.**
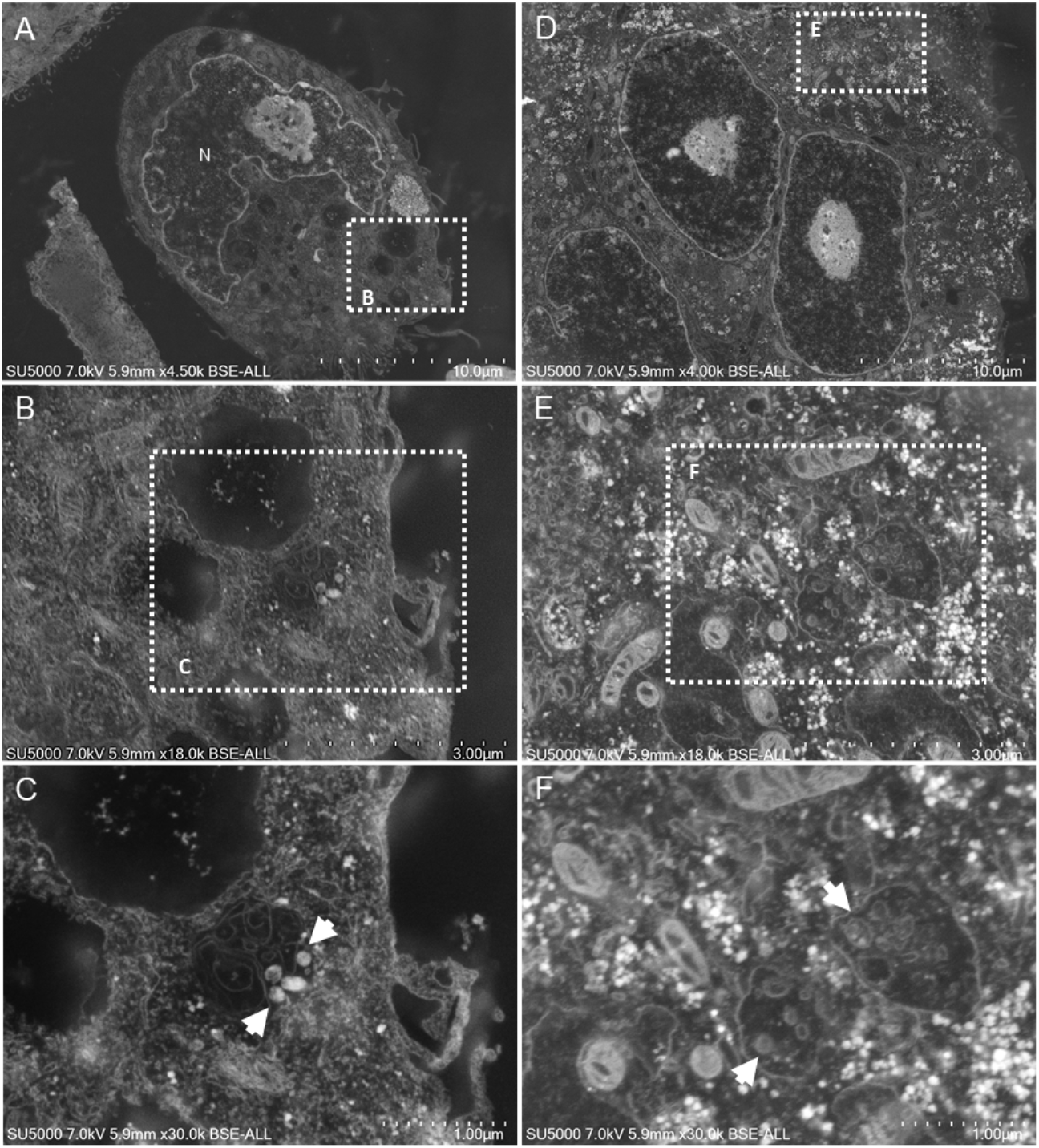
Scanning electron microscopy images of the ultra-thin section of a trophoblast cell incubated with *Coxiella* bacteria. **(A, B, C)** SEM images of *C. burnetii* Guiana strain infected BeWo cells. **(B)** High magnification of the boxed region in **(A)** shows serval vacuoles dispersed in the cell cytoplasm. **(C)** High magnification of the boxed region in **(B)** shows a vacuole containing electron-dense circular structures resembling *C. burnetii* (white arrows). **(D, E, F)** SEM images of *C. burnetii* Nine Mile strain infected BeWo cells. **(D)** High magnification of the boxed region in **(E)** shows vacuoles inside the BeWo cell **(F)** filled with hypo electron-dense *C. burnetii*-like structures with or without less electron-dense internal content (white arrows).

**Figure S2.**
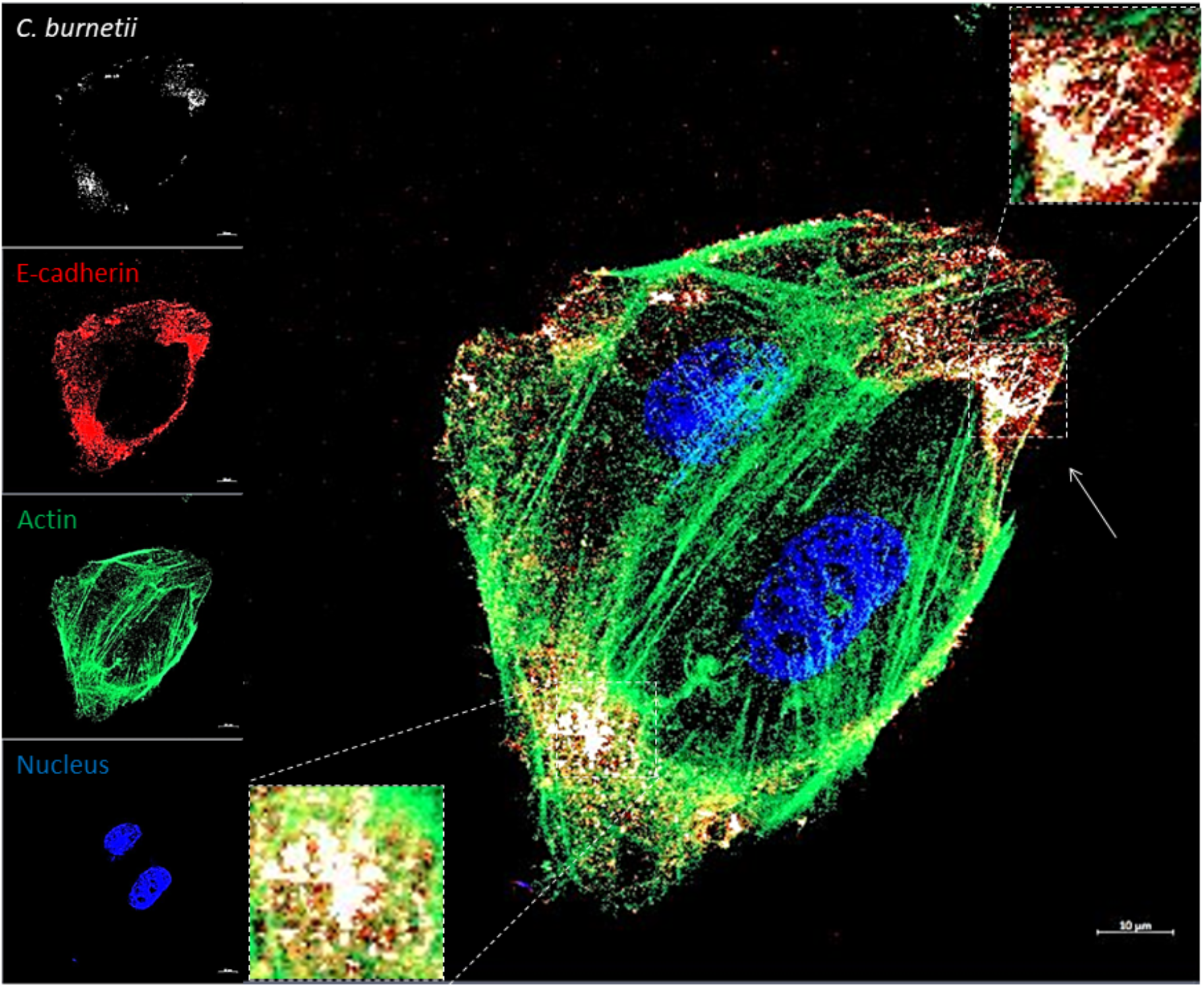
Confocal microscopy analysis of BeWo cells at 5 min post-infection with *C. burnetii*. Fluorescence labelling was as follows: *C. burnetii* (white), E-cad (red), actin (green), and cell nucleus (blue). Images were acquired using a confocal microscope (Zeiss LSM 800) with a 63X/1.4 oil objective (scale bar: 10 μm). Regions of high concentration of *C. burnetii* are shown with higher magnitude within scales.

**Figure S3.**
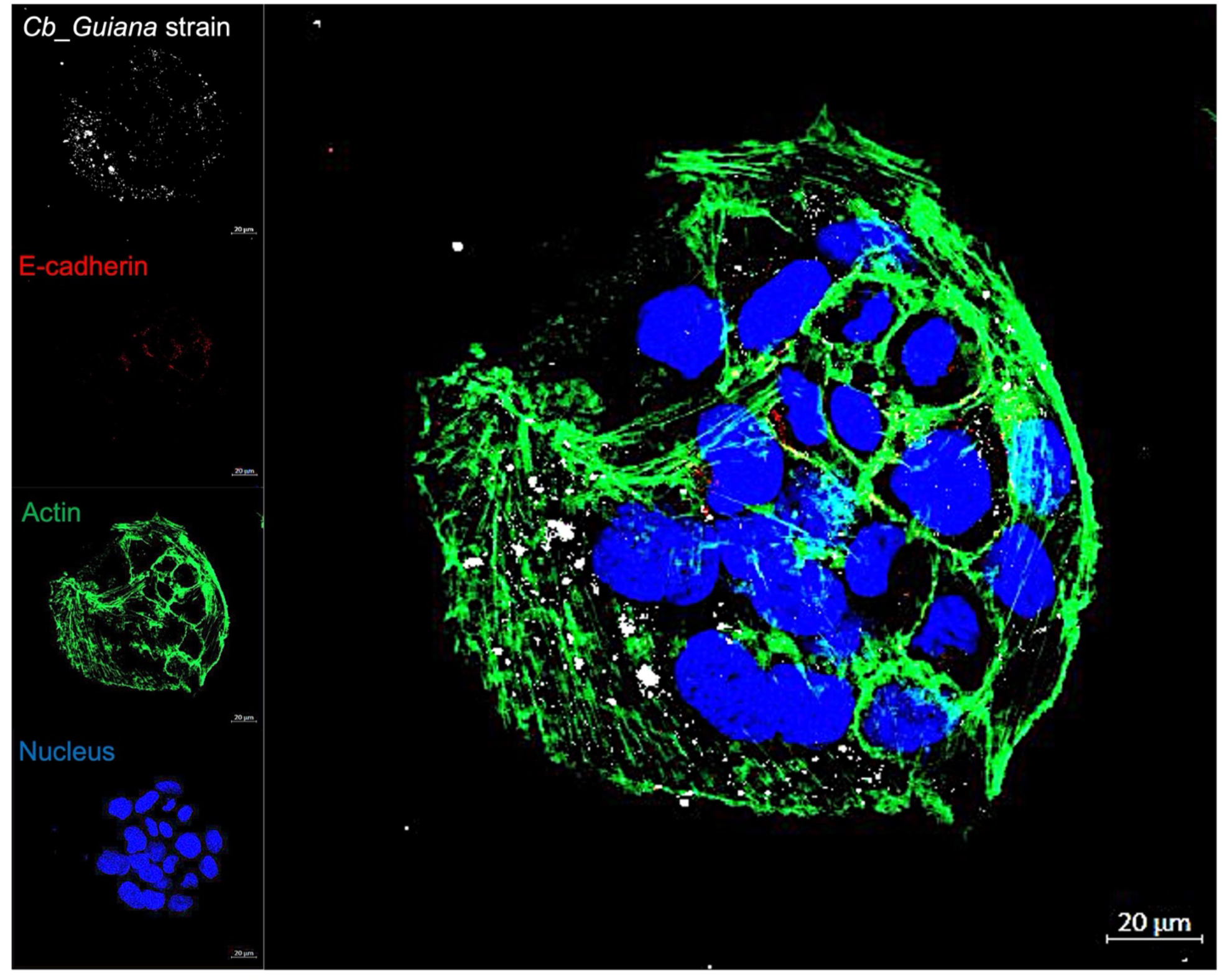
Confocal microscopy analysis of BeWo cells at 4 hours post-infection with *C. burnetii* (Cb) Guiana strain. Fluorescence labelling was as follows: *C. burnetii* (white), E-cad (red), actin (green), and cell nucleus (blue). Images were acquired using a confocal microscope (Zeiss LSM 800) with a 63X/1.4 oil objective (scale bar: 20 μm).

**Figure S4.**
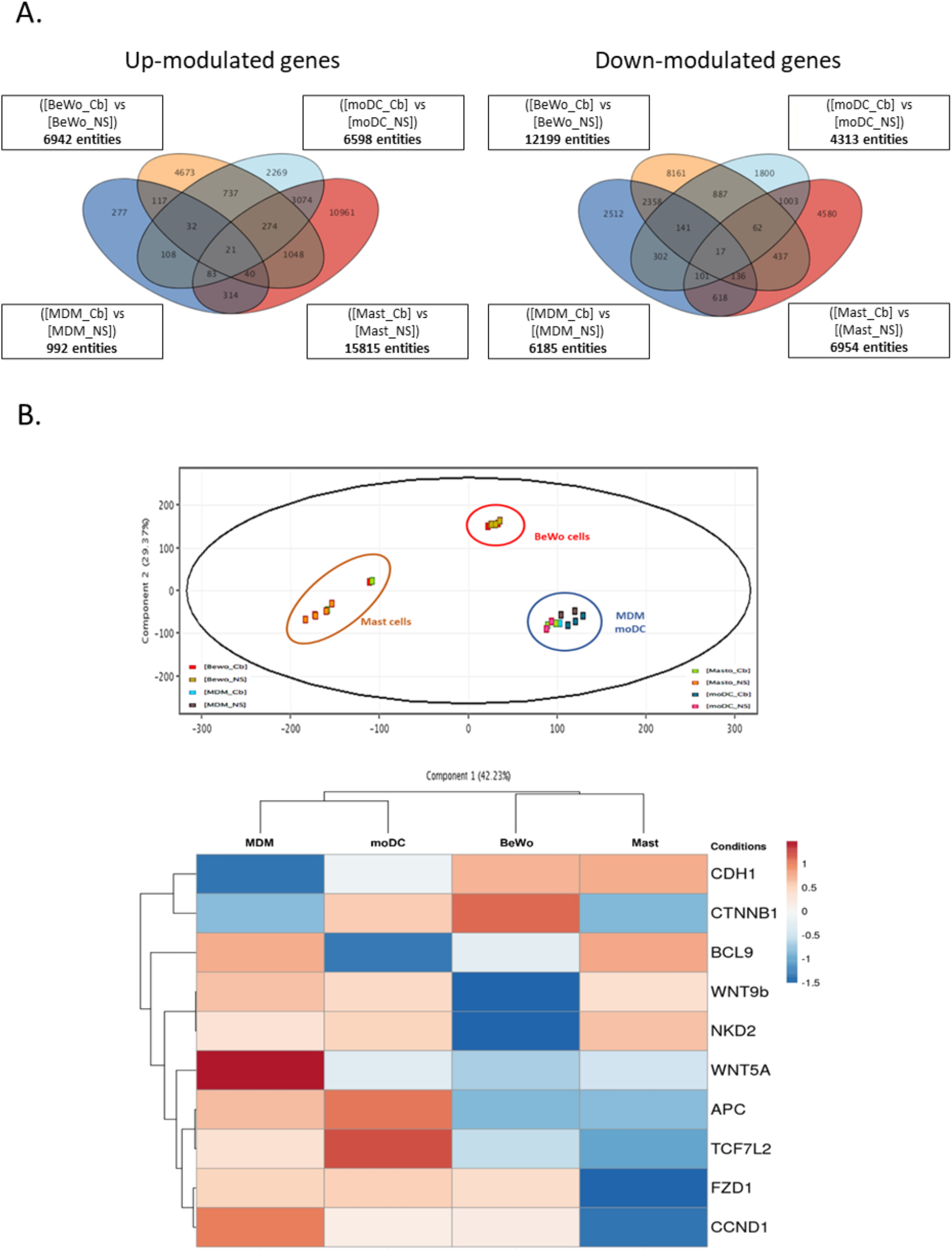
Exploration of gene signatures of *C. burnetii* infection (Nine Mile laboratory strain) of human cells as evidenced by microarray analysis. BeWo cells infected or not by *C. burnetii* were submitted to microarray analysis as well as primary cells, including monocyte-derived macrophages (MDM), monocyte-derived dendritic cells (moDC), and mast cells. **(A)** Set of genes up- or down-modulated in the panel of *C. burnetii* infected cells compared to uninfected cells (e.g., 6942 genes up-regulated and 12199 genes down-regulated in BeWo cells after *C. burnetii* infection). **(B)** principal component analysis and **(C)** hierarchical clustering were performed to show the distribution of the different groups of infected cells and the modulation of gene expression.

## Notes

### Competing Interest Statement

The authors have declared no competing interest.

### Summary of Updates

Title update

## References

1. Maurin M, Raoult D. Q fever. Clin Microbiol Rev. 1999;12:518–53.

2. Baca OG, Li Y-P, Kumar H. Survival of the Q fever agent Coxiella burnetii in the phagolysosome. Trends in Microbiology. 1994;2:476–80.

3. Ghigo E, Pretat L, Desnues B, Capo C, Raoult D, Mege J-L. Intracellular Life of Coxiella burnetii in Macrophages: An Update. Annals of the New York Academy of Sciences. 2009;1166:55–66.

4. Ben Amara A, Ghigo E, Le Priol Y, Lépolard C, Salcedo SP, Lemichez E, et al. Coxiella burnetii, the Agent of Q Fever, Replicates within Trophoblasts and Induces a Unique Transcriptional Response. PLoS One. 2010;5:e15315.

5. Bechah Y, Verneau J, Ben Amara A, Barry AO, Lépolard C, Achard V, et al. Persistence of Coxiella burnetii, the Agent of Q Fever, in Murine Adipose Tissue. PLoS One. 2014;9:e97503.

6. Raoult D, Marrie T, Mege J. Natural history and pathophysiology of Q fever. The Lancet Infectious Diseases. 2005;5:219–26.

7. van Schaik EJ, Chen C, Mertens K, Weber MM, Samuel JE. Molecular pathogenesis of the obligate intracellular bacterium Coxiella burnetii. Nat Rev Microbiol. 2013;11:561–73.

8. Melenotte C, Protopopescu C, Million M, Edouard S, Carrieri MP, Eldin C, et al. Clinical Features and Complications of Coxiella burnetii Infections From the French National Reference Center for Q Fever. JAMA Netw Open. 2018;1:e181580.

9. Thuny F, Grisoli D, Cautela J, Riberi A, Raoult D, Habib G. Infective Endocarditis: Prevention, Diagnosis, and Management. Canadian Journal of Cardiology. 2014;30:1046–57.

10. Martinez E, Cantet F, Fava L, Norville I, Bonazzi M. Identification of OmpA, a Coxiella burnetii protein involved in host cell invasion, by multi-phenotypic high-content screening. PLoS Pathog. 2014;10:e1004013.

11. Martinez E, Allombert J, Cantet F, Lakhani A, Yandrapalli N, Neyret A, et al. Coxiella burnetii effector CvpB modulates phosphoinositide metabolism for optimal vacuole development. Proc Natl Acad Sci U S A. 2016;113:E3260–3269.

12. Shipman M, Lubick K, Fouchard D, Gurram R, Grieco P, Jutila M, et al. Proteomic and Systems Biology Analysis of the Monocyte Response to Coxiella burnetii Infection. Wang T, éditeur. PLoS ONE. 2013;8:e69558.

13. Stead CM, Omsland A, Beare PA, Sandoz KM, Heinzen RA. Sec-mediated secretion by Coxiella burnetii. BMC Microbiol. 2013;13:222.

14. Vodkin MH, Williams JC, Stephenson EH. Genetic Heterogeneity among Isolates of Coxiella burnetii. Microbiology. 1986;132:455–63.

15. Hendrix LR, Samuel JE, Mallavia LP. Differentiation of Coxiella burnetii isolates by analysis of restriction-endonuclease-digested DNA separated by SDS-PAGE. Microbiology. 1991;137:269–76.

16. Jäger C, Willems H, Thiele D, Baljer G. Molecular characterization of Coxiella burnetii isolates. Epidemiol Infect. 1998;120:157–64.

17. Beare PA, Samuel JE, Howe D, Virtaneva K, Porcella SF, Heinzen RA. Genetic Diversity of the Q Fever Agent, Coxiella burnetii, Assessed by Microarray-Based Whole-Genome Comparisons. J Bacteriol. 2006;188:2309–24.

18. Hornstra HM, Priestley RA, Georgia SM, Kachur S, Birdsell DN, Hilsabeck R, et al. Rapid Typing of Coxiella burnetii. Li W, éditeur. PLoS ONE. 2011;6:e26201.

19. Hoover TA, Culp DW, Vodkin MH, Williams JC, Thompson HA. Chromosomal DNA Deletions Explain Phenotypic Characteristics of Two Antigenic Variants, Phase II and RSA 514 (Crazy), of the Coxiella burnetii Nine Mile Strain. Infect Immun. 2002;70:6726–33.

20. Toman R, Škultéty L. Structural study on a lipopolysaccharide from Coxiella burnetii strain Nine Mile in avirulent phase II. Carbohydrate Research. 1996;283:175–85.

21. Q FEVER: REPORT OF A CASE SIMULATING LYMPHOMA. Ann Intern Med. 1957;47:1030.

22. Melenotte C, Million M, Audoly G, Gorse A, Dutronc H, Roland G, et al. B-cell non-Hodgkin lymphoma linked to Coxiella burnetii. Blood. 2016;127:113–21.

23. van Roeden SE, van Houwelingen F, Donkers CMJ, Hogewoning SJ, de Lange Mma, van der Hoek W, et al. Exposure to Coxiella burnetii and risk of non-Hodgkin lymphoma: a retrospective population-based analysis in the Netherlands. The Lancet Haematology. 2018;5:e211–9.

24. Rothman N, Skibola CF, Wang SS, Morgan G, Lan Q, Smith MT, et al. Genetic variation in TNF and IL10 and risk of non-Hodgkin lymphoma: a report from the InterLymph Consortium. The Lancet Oncology. 2006;7:27–38.

25. Ghigo E, Capo C, Raoult D, Mege J-L. Interleukin-10 Stimulates Coxiella burnetii Replication in Human Monocytes through Tumor Necrosis Factor Down-Modulation: Role in Microbicidal Defect of Q Fever. Tuomanen EI, éditeur. Infect Immun. 2001;69:2345–52.

26. Melenotte C, Mezouar S, Ben Amara A, Benatti S, Chiaroni J, Devaux C, et al. A transcriptional signature associated with non-Hodgkin lymphoma in the blood of patients with Q fever. Dolcetti R, éditeur. PLoS ONE. 2019;14:e0217542.

27. Mezouar S, Omar Osman I, Melenotte C, Slimani C, Chartier C, Raoult D, et al. High Concentrations of Serum Soluble E-Cadherin in Patients With Q Fever. Front Cell Infect Microbiol. 2019;9:219.

28. van Roeden SE, van Houwelingen F, Donkers CMJ, Hogewoning SJ, de Lange Mma, van der Hoek W, et al. Exposure to Coxiella burnetii and risk of non-Hodgkin lymphoma: a retrospective population-based analysis in the Netherlands. Lancet Haematol. 2018;5:e211–9.

29. Weehuizen JM, van Roeden SE, Hogewoning SJ, van der Hoek W, Bonten MJM, Hoepelman AIM, et al. No increased risk of mature B-cell non-Hodgkin lymphoma after Q fever detected: results from a 16-year ecological analysis of the Dutch population incorporating the 2007-2010 Q fever outbreak. Int J Epidemiol. 2022;dyac053.

30. Dash S, Duraivelan K, Samanta D. Cadherin-mediated host–pathogen interactions. Cellular Microbiology [Internet]. 2021 [cité 18 mars 2022];23. Disponible sur: https://onlinelibrary.wiley.com/doi/10.1111/cmi.13316

31. Ashton-Key M, Cowley GP, Smith ME. Cadherins in reactive lymph nodes and lymphomas: high expression in anaplastic large cell lymphomas. Histopathology. 1996;28:55–9.

32. McCrea PD, Maher MT, Gottardi CJ. Nuclear Signaling from Cadherin Adhesion Complexes. aCurrent Topics in Developmental Biology [Internet]. Elsevier; 2015 [cité 23 mars 2022]. p. 129–96. Disponible sur: https://linkinghub.elsevier.com/retrieve/pii/S0070215314000192

33. Vleminckx K, Vakaet L, Mareel M, Fiers W, Van Roy F. Genetic manipulation of E-cadherin expression by epithelial tumor cells reveals an invasion suppressor role. ell. 1991;66:107–19.

34. Christofori G, Semb H. The role of the cell-adhesion molecule E-cadherin as a tumour-suppressor gene. Trends in Biochemical Sciences. 1999;24:73–6.

35. Pe N. Tumor suppressor gene E-cadherin and its role in normal and malignant cells. Cancer Cell International. 2003;7.

36. Wells A, Yates C, Shepard CR. E-cadherin as an indicator of mesenchymal to epithelial reverting transitions during the metastatic seeding of disseminated carcinomas. Clin Exp Metastasis. 2008;25:621–8.

37. Izaguirre MF, Galetto CD, Baró L, Casco VH. E-Cadherin Dysfunction and Cancer. Journal of Biosciences and Medicines. Scientific Research Publishing; 2019;7:42–67.

38. Na T-Y, Schecterson L, Mendonsa AM, Gumbiner BM. The functional activity of E-cadherin controls tumor cell metastasis at multiple steps. Proc Natl Acad Sci USA. 2020;117:5931–7.

39. Grabowska MM, Day ML. Soluble E-cadherin: More Than a Symptom of Disease. Front Biosci (Landmark Ed). 2012;17:1948–64.

40. Devaux CA, Mezouar S, Mege J-L. The E-Cadherin Cleavage Associated to Pathogenic Bacteria Infections Can Favor Bacterial Invasion and Transmigration, Dysregulation of the Immune Response and Cancer Induction in Humans. Front Microbiol. 2019;10:2598.

41. Aghababaei M, Hogg K, Perdu S, Robinson WP, Beristain AG. ADAM12-directed ectodomain shedding of E-cadherin potentiates trophoblast fusion. Cell Death Differ. 2015;22:1970–84.

42. Capo C, Lindberg FP, Meconi S, Zaffran Y, Tardei G, Brown EJ, et al. Subversion of monocyte functions by coxiella burnetii: impairment of the cross-talk between alphavbeta3 integrin and CR3. J Immunol. 1999;163:6078–85.

43. Ghigo E, Imbert G, Capo C, Raoult D, Mege J-L. Interleukin-4 Induces Coxiella burnetii Replication in Human Monocytes but not in Macrophages. Annals of the New York Academy of Sciences. 2003;990:450–9.

44. Heystek HC, Mudde GC, Ohler R, Kalthoff FS. Granulocyte-macrophage colony-stimulating factor (GM-CSF) has opposing effects on the capacity of monocytes versus monocyte-derived dendritic cells to stimulate the antigen-specific proliferation of a human T cell clone. Clin Exp Immunol. 2000;120:440–7.

45. Saito H, Kato A, Matsumoto K, Okayama Y. Culture of human mast cells from peripheral blood progenitors. Nat Protoc. 2006;1:2178–83.

46. Winer J, Jung CKS, Shackel I, Williams PM. Development and Validation of Real-Time Quantitative Reverse Transcriptase–Polymerase Chain Reaction for Monitoring Gene Expression in Cardiac Myocytesin Vitro. Analytical Biochemistry. 1999;270:41–9.

47. Shannon JG, Heinzen RA. Adaptive immunity to the obligate intracellular pathogen Coxiella burnetii. Immunol Res. 2009;43:138–48.

48. Sireci G, Badami GD, Di Liberto D, Blanda V, Grippi F, Di Paola L, et al. Recent Advances on the Innate Immune Response to Coxiella burnetii. Front Cell Infect Microbiol. 2021;11:754455.

49. Zarza SM, Mezouar S, Mege J-L. From Coxiella burnetii Infection to Pregnancy Complications: Key Role of the Immune Response of Placental Cells. Pathogens. 2021;10:627.

50. Graham JG, Winchell CG, Kurten RC, Voth DE. Development of an Ex Vivo Tissue Platform To Study the Human Lung Response to Coxiella burnetii. Roy CR, éditeur. Infect Immun. 2016;84:1438–45.

51. Dragan AL, Kurten RC, Voth DE. Characterization of Early Stages of Human Alveolar Infection by the Q Fever Agent Coxiella burnetii. Roy CR, éditeur. Infect Immun. 2019;87:e00028–19.

52. Coutifaris C, Kao LC, Sehdev HM, Chin U, Babalola GO, Blaschuk OW, et al. E-cadherin expression during the differentiation of human trophoblasts. Development. 1991;113:767–77.

53. Lecuit M, Nelson DM, Smith SD, Khun H, Huerre M, Vacher-Lavenu M-C, et al. Targeting and crossing of the human maternofetal barrier by Listeria monocytogenes : Role of internalin interaction with trophoblast E-cadherin. Proc Natl Acad Sci USA. 2004;101:6152–7.

54. Coleman SA, Fischer ER, Howe D, Mead DJ, Heinzen RA. Temporal Analysis of Coxiella burnetii Morphological Differentiation. J Bacteriol. 2004;186:7344–52.

55. Meconi S, Capo C, Remacle-Bonnet M, Pommier G, Raoult D, Mege J-L. Activation of Protein Tyrosine Kinases by Coxiella burnetii: Role in Actin Cytoskeleton Reorganization and Bacterial Phagocytosis. Infect Immun. 2001;69:2520–6.

56. Voth DE, Heinzen RA. Sustained Activation of Akt and Erk1/2 Is Required for Coxiella burnetii Antiapoptotic Activity. Infect Immun. 2009;77:205–13.

57. Zheng G, Lyons JG, Tan TK, Wang Y, Hsu T-T, Min D, et al. Disruption of E-Cadherin by Matrix Metalloproteinase Directly Mediates Epithelial-Mesenchymal Transition Downstream of Transforming Growth Factor-β1 in Renal Tubular Epithelial Cells. The American Journal of Pathology. 2009;175:580–91.

58. Marambaud P. A presenilin-1/gamma-secretase cleavage releases the E-cadherin intracellular domain and regulates disassembly of adherens junctions. The EMBO Journal. 2002;21:1948–56.

59. Kourtidis A, Ngok SP, Anastasiadis PZ. p120 Catenin. Progress in Molecular Biology and Translational Science [Internet]. Elsevier; 2013 [cité 23 mars 2022]. p. 409–32. Disponible sur: https://linkinghub.elsevier.com/retrieve/pii/B9780123943118000182

60. O’Connor PM, Lapointe TK, Jackson S, Beck PL, Jones NL, Buret AG. Helicobacter pylori Activates Calpain via Toll-Like Receptor 2 To Disrupt Adherens Junctions in Human Gastric Epithelial Cells. Blanke SR, éditeur. Infect Immun. 2011;79:3887–94.

61. Wu S, Morin PJ, Maouyo D, Sears CL. Bacteroides fragilis enterotoxin induces c-Myc expression and cellular proliferation. Gastroenterology. 2003;124:392–400.

62. Chung L, Thiele Orberg E, Geis AL, Chan JL, Fu K, DeStefano Shields CE, et al. Bacteroides fragilis Toxin Coordinates a Pro-carcinogenic Inflammatory Cascade via Targeting of Colonic Epithelial Cells. Cell Host & Microbe. 2018;23:203-214.e5.

63. Meconi S, Jacomo V, Boquet P, Raoult D, Mege J-L, Capo C. Coxiella burnetii Induces Reorganization of the Actin Cytoskeleton in Human Monocytes. Infect Immun. 1998;66:5527–33.

64. Ayollo DV, Zhitnyak IY, Vasiliev JM, Gloushankova NA. Rearrangements of the Actin Cytoskeleton and E-Cadherin–Based Adherens Junctions Caused by Neoplasic Transformation Change Cell–Cell Interactions. Hartl D, éditeur. PLoS ONE. 2009;4:e8027.

65. Khan NI, Bradstock KF, Bendall LJ. Activation of Wnt/?-catenin pathway mediates growth and survival in B-cell progenitor acute lymphoblastic leukaemia. Br J Haematol. 2007;138:338–48.

66. Mani M, Carrasco DE, Zhang Y, Takada K, Gatt ME, Dutta-Simmons J, et al. BCL9 Promotes Tumor Progression by Conferring Enhanced Proliferative, Metastatic, and Angiogenic Properties to Cancer Cells. Cancer Res. 2009;69:7577–86.

67. Osman IO, Caputo A, Pinault L, Mege J-L, Levasseur A, Devaux CA. Identification and characterization of an HtrA sheddase produced by Coxiella burnetii [Internet]. bioRxiv; 2023 [cité 10 févr 2023]. p. 2023.01.26.525556. Disponible sur: https://www.biorxiv.org/content/10.1101/2023.01.26.525556v1

